# nf-core/airrflow: an adaptive immune receptor repertoire analysis workflow employing the Immcantation framework

**DOI:** 10.1101/2024.01.18.576147

**Authors:** Gisela Gabernet, Susanna Marquez, Robert Bjornson, Alexander Peltzer, Hailong Meng, Edel Aron, Noah Y. Lee, Cole Jensen, David Ladd, Friederike Hanssen, Simon Heumos, nf-core community, Gur Yaari, Markus C. Kowarik, Sven Nahnsen, Steven H. Kleinstein

**Author notes:** Equal contribution (Gisela Gabernet, Susanna Marquez). co-senior authors (Sven Nahnsen, Steven H. Kleinstein).

## Abstract

Adaptive Immune Receptor Repertoire sequencing (AIRR-seq) is a valuable experimental tool to study the immune state in health and following immune challenges such as infectious diseases, (auto)immune diseases, and cancer. Several tools have been developed to reconstruct B cell and T cell receptor sequences from AIRR-seq data and infer B and T cell clonal relationships. However, currently available tools offer limited parallelization across samples, scalability or portability to high-performance computing infrastructures. To address this need, we developed nf-core/airrflow, an end-to-end bulk and single-cell AIRR-seq processing workflow which integrates the Immcantation Framework following BCR and TCR sequencing data analysis best practices. The Immcantation Framework is a comprehensive toolset, which allows the processing of bulk and single-cell AIRR-seq data from raw read processing to clonal inference. nf-core/airrflow is written in Nextflow and is part of the nf-core project, which collects community contributed and curated Nextflow workflows for a wide variety of analysis tasks. We assessed the performance of nf-core/airrflow on simulated sequencing data with sequencing errors and show example results with real datasets. To demonstrate the applicability of nf-core/airrflow to the high-throughput processing of large AIRR-seq datasets, we validated and extended previously reported findings of convergent antibody responses to SARS-CoV-2 by analyzing 97 COVID-19 infected individuals and 99 healthy controls, including a mixture of bulk and single-cell sequencing datasets. Using this dataset, we extended the convergence findings to 20 additional subjects, highlighting the applicability of nf-core/airrflow to validate findings in small in-house cohorts with reanalysis of large publicly available AIRR datasets.

**Availability and implementation:** nf-core/airrflow is available free of charge, under the MIT license on GitHub (https://github.com/nf-core/airrflow). Detailed documentation and example results are available on the nf-core website at (https://nf-co.re/airrflow).

**Visual abstract:** 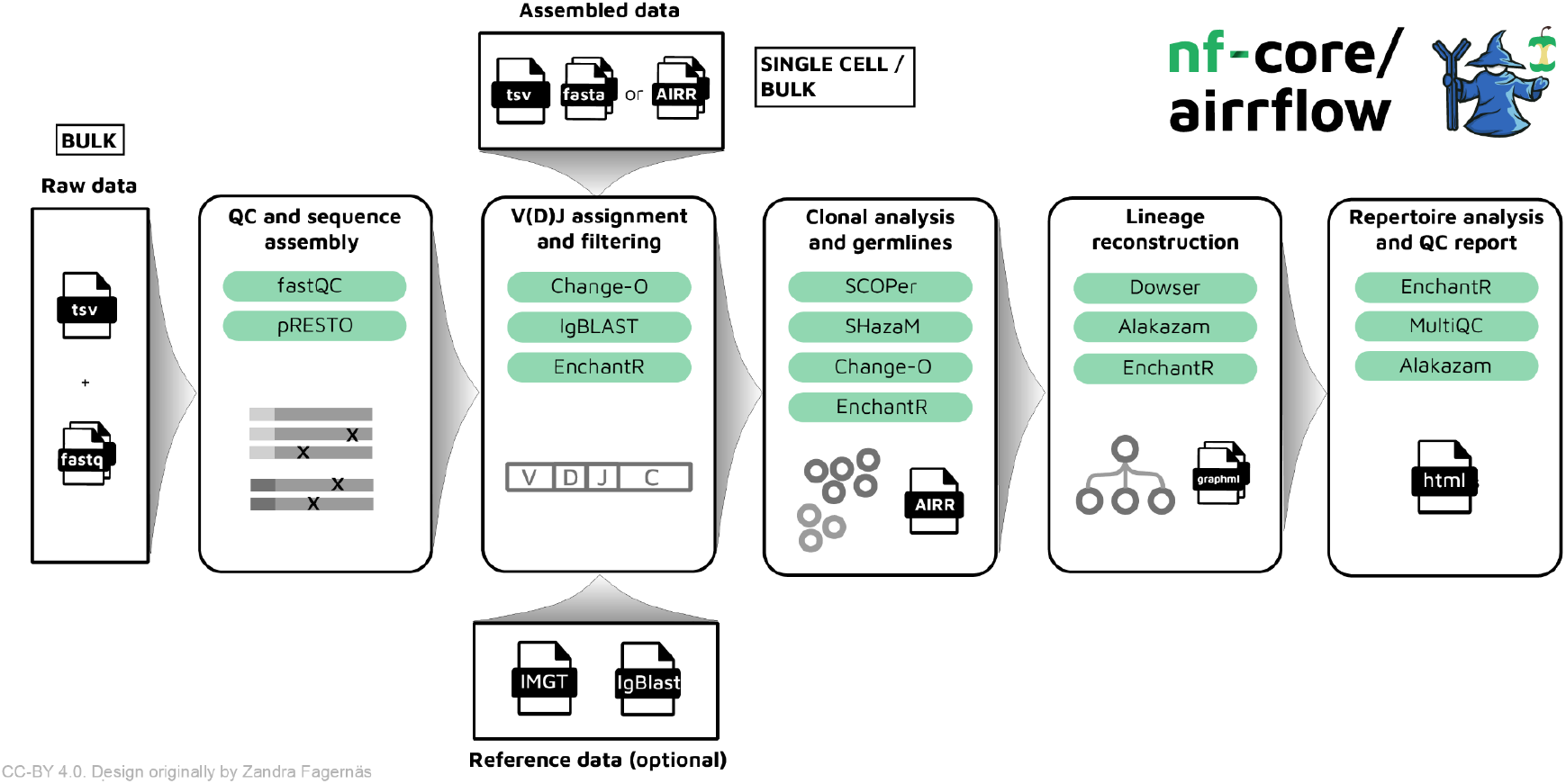

## Introduction

B cells and T cells, key components of adaptive immunity, recognize foreign pathogens and provide long-term protection against them. They are also implicated in auto-immune diseases when eliciting a deleterious response against self-antigens. The antigen recognition is performed through membrane receptors, termed B cell and T cell receptors (BCR and TCR, respectively). BCRs in their secreted form are termed antibodies. BCRs and TCRs are generated during cell maturation by the somatic DNA recombination of a number of variable (V), diversity (D), and joining (J) gene segments in the immunoglobulin (IGH, IGK and IGL) and TCR (TRA and TRB) loci(1,2). Additional nucleotide deletion/insertion at the gene boundaries generate practically unique receptors for each maturing cell(3,4). The constant region is encoded in additional exons and added to the sequence during transcription and splicing. Two identical heavy and light chains constitute the BCR, while an alpha and a beta chain constitute the TCR. In BCRs and antibodies, the heavy chain constant region determines the isotype. The diversity of BCRs is further increased upon antigen encounter and activation by a process termed somatic hypermutation (SHM), which provides the substrate for antigen driven selection leading to increased affinity to their targets(5). The collection of BCRs and TCRs in an individual, tissue, or cell subset is referred to as the adaptive immune receptor repertoire (AIRR). Characterizing AIRRs is relevant for the study of the immune state of individuals in health and following alterations such as vaccines(6,7), infectious diseases(8,9), (auto)immune diseases(10–13), and cancer(14).

Adaptive Immune Receptor Repertoire sequencing (AIRR-seq) enables the recovery of the collection of AIRRs sampled from an individual(15). AIRR-seq can be performed from the genomic DNA or expressed RNA libraries. Methods based on the targeted amplification of the expressed BCR and TCR transcripts are more widely used due to the possibility of additionally sequencing the constant region, which determines the BCR isotype, and the presence of multiple RNA molecules per cell that can enhance sensitivity and allow for error correction when employing protocols that incorporate Unique Molecular Identifiers (UMIs). Single-cell sequencing technologies have revolutionized the field by allowing the characterization of the paired heavy and light (BCR) or alpha and beta (TCR) chain receptor sequences linked to the individual cell’s transcriptomic profiles(16). The processing of AIRR-seq data requires steps that are distinct from other bioinformatics pipelines(17,18), which has led to the development of a multitude of tools and frameworks(11,19–33). However, individual tools offer limited parallelization across samples, scalability or native support for alternative compute infrastructures such as high-performance computing (HPC) clusters or commercial clouds.

Workflow management systems such as Nextflow(34) or Snakemake(35) allow the incorporation of tools into highly configurable analysis workflows which can be triggered with a single command, and offer implicit massive parallelization across the analyzed samples. Workflow management systems additionally help ensure reproducible analysis results in several ways: by stating the commands used to execute the individual tools, reporting the parameters used to launch the workflow, and easing the use of container engines in the individual analysis steps. As an example, Nextflow supports the download and execution of Docker, Apptainer, and other container engines from public container repositories by solely specifying their registry address, name and version in the desired workflow steps. Volume mounts for the start and stop commands are set by Nextflow in the background. Containers provide a controlled running environment with fixed tool versions and system libraries, which is critical in ensuring reproducible results and portability across compute infrastructures, such as clouds(34).

To address the need for a high-throughput, portable analysis workflow which incorporates the necessary steps for AIRR-seq data analysis we developed nf-core/airrflow, a flexible workflow written in the Nextflow language. The workflow utilizes the Immcantation framework, a comprehensive collection of open-source software to process AIRR-seq data from start-to-finish, covering sequence assembly and error correction(19), alignment to an international immunogenetics information system (IMGT)(36,37) BCR and TCR reference data with IgBLAST(38), clonal relationship inference(20,21,25,39), reconstruction of clonal lineages(23,24), and the identification of predictive repertoire properties and sequence motifs(11,40). nf-core/airrflow is a flexible workflow which supports the analysis of both bulk and single-cell sequencing datasets generated with a multitude of protocols, allowing as well the combined processing of datasets with mixed bulk and single-cell modalities. The workflow is part of the nf-core project(41), which defines Nextflow implementation best practices including containerization of all software tools with biocontainers(42) whenever possible, reporting of all software versions in a MultiQC(43) report, continuous integration testing with example test data, and portability testing to cloud infrastructures. These features aim at making nf-core/airrflow easy to use and install, requiring only Nextflow and a container engine as dependencies, as well as ensuring that the obtained results will be reproducible and portable across compute infrastructures.

We benchmarked the workflow on simulated ground truth BCR repertoire sequencing data and showed its superiority to alternative existing tools. As an application use-case, we validated the findings of convergent antibody sequences across three COVID-19 diagnosed individuals identified by Robbiani *et al*.(44) in a larger dataset comprising BCR repertoire samples from 97 individuals diagnosed with COVID-19 and 99 healthy controls retrieved from the from the AIRR Data Commons(45) by querying the iReceptor Gateway(46). Thus, we showcased the applicability of nf-core/airrflow to validate findings in small patient cohorts with data from publicly available sources.

## Design and implementation

### Workflow outline

nf-core/airrflow is a high-throughput workflow for the end-to-end analysis of AIRR bulk and single-cell sequencing data utilizing the Immcantation framework(11,19–25). It encompasses sequencing read assembly, V(D)J and constant region allele assignment, clonal inference, and repertoire analysis including quality filtering and quality control reporting in each step (Figure 1A). Multiple protocols for raw bulk sequencing data analysis are supported, including multiplexed PCR and 5’ Rapid amplification of cDNA ends (5’-RACE) based protocols, with or without the inclusion of UMIs that allow the correction of sequencing errors and amplification biases. The analysis of assembled bulk and single-cell BCR and TCR sequencing data generated with the 10x genomics platform or other platforms provided in the AIRR rearrangement format is also supported(47). The different options can be specified through command line parameters when executing a pipeline or a Nextflow parameters file that is provided at runtime.

**Figure 1.**
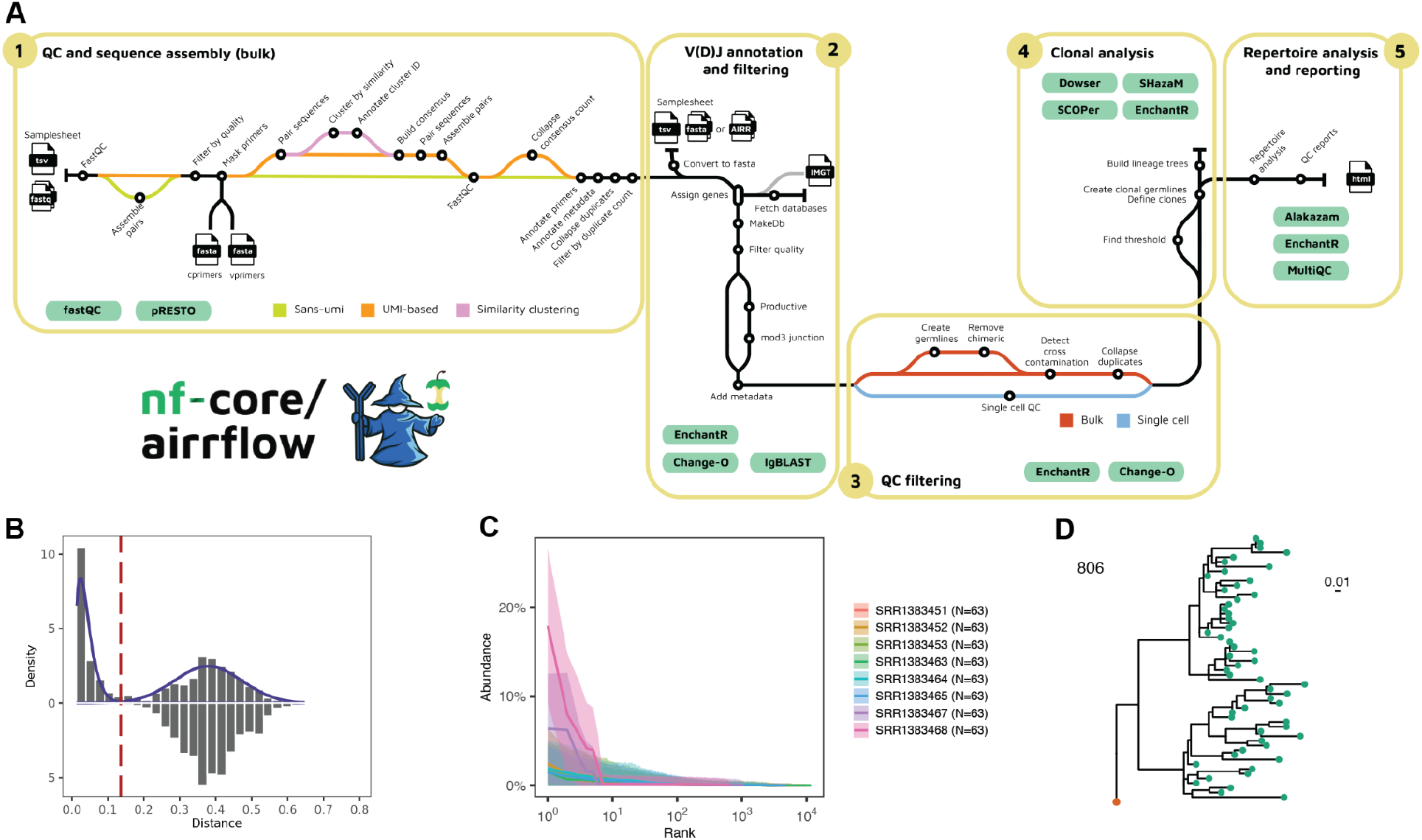
Schematic representation of the nf-core/airrflow workflow processes and detailed analysis steps. **A**. Workflow overview. QC and sequence assembly steps (1) are performed on the bulk raw sequencing reads. The workflow supports several sequencing protocols including multiplex PCR and 5’-RACE, with or without UMIs. For single-cell sequencing data or readily-assembled bulk sequencing reads, the analysis starts with V(D)J and C gene assignment with IgBLAST (2). Reference BCR and TCR sequences can directly be pulled from IMGT, or provided by the user for enhanced reproducibility. After V(D)J assignment, the sequences are filtered for meaningful alignments (minimum of base alignments, as well as % of accepted N nucleotides) and productive sequences. Additional QC steps can be applied to remove chimeric reads and duplicate sequences, and detect cross-sample contamination (3). Clonal groups are defined by hierarchical clustering using a hamming distance threshold which can be provided or estimated from the data (4). Lineage trees can be optionally reconstructed with the Dowser package. A final report summarizes the repertoire analysis results (5). MultiQC integrates the quality control results of the QC steps and reports the software versions of all executed processes in the workflow. To generate example data, nf-core/airrflow was run on full-size test data on AWS batch extracted from the publication of Stern et al.(11) Example of a hamming distance threshold plot (**B**), clonal abundance plot (**C**), and reconstructed lineage tree (**D**) from this dataset.

### Input data

The bulk raw AIRR-seq data processing starts from the FastQ files. Those are specified within a metadata file following the AIRR Study Reporting standard (MiAIRR)(48), which includes the sample identifiers, species, target locus (IG or TR), and other information required for the repertoire analysis. The primer and/or RACE linker sequences used for library preparation are additional required inputs for processing bulk AIRR-seq data. Examples of supported kits are the commercially available New England Biolabs kits(49), or TAKARA Bio BCR and TCR profiling kits(50). It is possible to specify in a flexible manner the start position of the primer(s) and UMI position and length, extending the applicability to other custom protocols. For single-cell AIRR-seq data analysis, the pipeline accepts as input sequence tables in the AIRR rearrangement format(48), such as the ones provided by the 10X Genomics Cell Ranger, or tools that can extract BCR and TCR sequences from single-cell RNA-seq sequencing data such as TRUST4(51). The analysis workflow then starts with the optional V(D)J and constant region allele reassignment with IgBLAST, to ensure that the sequences are annotated with the same IMGT(36) BCR and TCR reference data, and IMGT gaps are included in the alignments. It is also possible to process readily assembled bulk sequencing data starting with this step, by providing the fully assembled and error-corrected reads in fasta or AIRR rearrangement format.

### Read QC and sequence assembly

Sequencing read quality control is performed using fastp(52) for general read quality statistics and adapter trimming, if indicated. The pRESTO(19) Immcantation tool is employed for all other sequence assembly and error correction steps. Pre-processing involves the filtering of low-quality reads, extraction of UMIs, and primer masking. For protocols including UMIs, a consensus sequence is built with sequences comprising the same UMI. Optionally, the *cluster set* process allows dealing with insufficient UMI diversity, which might occur when UMIs are too short for the library diversity. Read mates are paired and assembled, and the primer information along with any metadata relevant to the downstream analysis is annotated in the fastq header with a format standardized by pRESTO. Duplicate sequences are collapsed, annotating the individual number of representative reads where this sequence was observed. Finally, sequences with insufficient support, less than two representatives by default, are filtered out.

### V(D)J annotation and alignment QC

V(D)J and constant region allele assignment is performed by aligning the assembled reads to the relevant BCR and TCR reference data with IgBLAST(37,38), and processing the alignment results with the Change-O(20) Immcantation tool. Currently, human and mouse IMGT reference data are supported and directly downloaded by the workflow. Alternatively, users can provide curated custom reference data for other species in the same format. Several QC and filtering steps are performed after alignment. First, the provided desired locus is checked to match the assigned V gene chain. By default, sequence alignments that contain less than 200 informative positions, more than 10% N nucleotides, or that generate non-productive transcripts are filtered out. For bulk sequencing data, chimeric reads sometimes generated during PCR amplification can be optionally removed. Likely cross-sample sequence contamination is detected by identifying samples with a high percentage of overlapping identical VDJ sequences. As a last step for bulk sequencing QC, duplicated sequences are collapsed and the duplicate number is annotated. Filtering and quality control steps for single-cell sequencing data include removing cells containing multiple heavy chains or only light chains, as well as detecting and removing cross-sample sequence contaminants with identical VDJ region sequences and cell barcodes.

### Clonal inference

Clonal inference involves clustering the set of BCR and TCR sequences into clones, which are defined as the group of cells that descend from a common ancestor. This is a harder problem for B cells than T cells, as SHM introduces targeted mutations in the BCR sequences during B cell clonal expansion, generating a diversity of sequences within the same clone that are relevant for antibody affinity maturation. Clonal relationships are identified with the Immcantation SCOPer(21,22) integrated methods. By default, sequences are partitioned into groups with the same V gene (IGHV or TRBV), J gene (IGHJ or TRBJ) and junction length - the junction sequence corresponds to the highly variable complementarity determining region 3 (CDR3) region with additional flanking nucleotides that encode two conserved amino acids in the 5’ and 3’ end -. Hierarchical clustering with single-linkage is applied to the heavy chain junction nucleotide sequence, with the length-normalized hamming distance as a distance metric. For B cell repertoires, the hamming distance threshold used to distinguish sequences belonging to the same clonal group can be estimated using the SHazaM(53,54) Immcantation package. As T cells do not undergo somatic hypermutation, T cell clones are defined as having identical TCR sequences. In the current implementation, TCR sequences sharing the same TRBV and TRBJ alleles and displaying identical junction nucleotide sequences (hamming distance threshold of zero) are assigned to the same clone, because in some sequencing protocols the full V(D)J sequence will not be recovered. For single-cell data where paired heavy and light chains are available, the clones can be additionally partitioned according to the light chain (or alpha chain) V genes, J genes and junction length. Summary reports of the identified distance threshold for all the samples and clonal relationships facilitate a visual validation of the results (Figure 1B). Clonal repertoire properties such as clonal abundance and diversity are calculated with the Alakazam Immcantation package (Figure 1C). Clonal lineage trees can be optionally reconstructed with Dowser(55), an Immcantation tool which includes several methods for lineage tree reconstruction such as IgPhyML(23), a method specifically designed for B-cell lineage tree reconstruction (Figure 1D).

### Reporting

HTML reports on individual workflow steps can be found in the respective process output folders, and are generated through EnchantR, an R helper package developed along with the Nextflow workflow, which is now part of Immcantation. An interactive report is generated comprising the number of sequences obtained after each workflow step. The report in Rmarkdown format can be downloaded by the user and modified accordingly to meet the specific project analysis needs. Additionally, a MultiQC(43) interactive report is generated, which contains a read quality control report over all the processed samples and the software versions employed by each workflow process.

## Results

### nf-core/airrflow benchmarking with simulated data

To evaluate the ability of nf-core/airrflow to recover immune repertoire sequences and infer clonal relationships from ground truth sequencing data, we simulated three BCR receptor repertoires with known V(D)J sequences, varying clonal abundances and increasing frequency of sequencing errors. The germline sequences were generated by simulating V(D)J recombination with ImmuneSIM(56). Clonal lineage trees and somatic hypermutation were simulated with SHazaM to obtain a power law and a uniform clonal size distribution (repA and repB, respectively) or extracted from a real BCR repertoire sample previously published(57) (repC). 5000 singleton sequences - representing naive B cells that are not clonally expanded - were added to the synthetic repertoires repA and repB to achieve a similar frequency as observed in the real BCR repertoire repC (see Supplementary methods and Supplementary Figure 1 for further details on the repertoire simulation).

We then simulated paired-end raw sequencing data for each repertoire with Grinder(58). To assess the impact of sequencing errors on the final sequence recovery, the sequencing data simulations were performed with increasing percentages of sequencing errors modeled in a linear fashion along the read length, increasing from zero to a predetermined percentage, in accordance with previous studies on Illumina sequencing error values and their distribution along the read positions(59,60). Five simulated libraries were prepared, with 0%, 0.1%, 0.25%, 0.5%, 1.0% sequencing errors in the center of the reads. Additionally, two library preparation protocols were compared: with (*UMI*) and without (*sans-UMI*) UMIs. Adding UMIs to the library preparation procedure offers the potential for sequencing error correction by constructing a consensus sequence of the recovered sequences with identical UMIs, which is a widely used strategy in BCR and TCR sequencing protocols.

The simulated BCR sequencing libraries were processed with nf-core/airrflow, and the ability to recover the original sequences in the simulated repertoires was evaluated (Figure 2, Suppl. Figure 2). We evaluated the proportion of correctly identified sequences for each of the repertoires with exact sequence matches (sensitivity exact matches) and additionally considered the matches of sequences that contain “N” nucleotides (Figure 2A and B) for a protocol including UMIs. The N-nucleotides are introduced when UMIs are used for error correction by building a sequence consensus, and the consensus base is under a certain frequency threshold (default minimum frequency 0.6) or quality threshold (default minimum quality 0) so there is insufficient consensus to call a particular base. nf-core/airrflow correctly recovered over 99% of the sequences in each of the three repertoires when no simulated sequencing errors were present. The data with sequencing errors decreased the proportion of correctly identified sequences, but sensitivity was maintained above 97% for exact sequence matches and 98% for matches containing “N” nucleotides due to not reaching sufficient consensus. Two incorrect sequences were reported due to a rare occurrence of duplicate UMIs being assigned to two highly similar sequences that are part of the same clone, and two sequences with the same UMI with simulated errors at the same position (Figure 2 C). The number of missing sequences ranged from 100 to 300 from a total of 21,321 (repA), 20,959 (repB), and 15,329 (repC). The sensitivity and number of missing sequences was comparable to MiXCR(26), an alternative tool for bulk and single-cell AIRR-seq data analysis (Figure 2 A-D, Supplementary Table 1). When simulating a protocol without UMI, the sensitivity started at 99% with the absence of sequencing errors, but dropped below 50% and up to 125 incorrect sequences with increasing sequencing errors (Suppl. Figure 3A and B), reinforcing the importance of utilizing protocols that include UMI error correction. The high number of missing sequences is due to a quality control step that eliminates sequences that do not contain at least two representative copies, which are attributed to sequencing errors (Suppl. Figure 3C).

**Figure 2.**
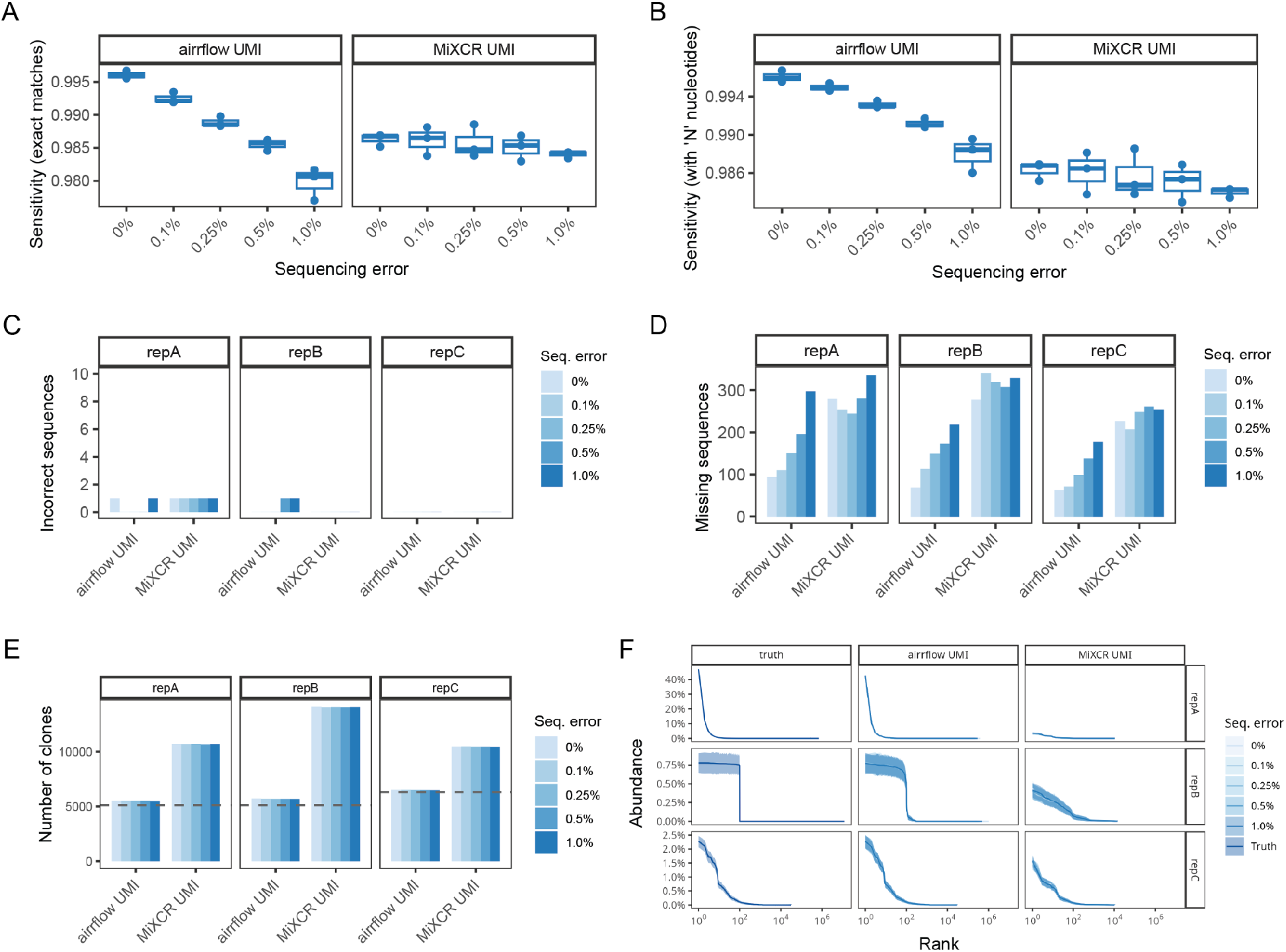
Performance assessment of the nf-core/airrflow pipeline on three simulated BCR repertoires compared to MiXCR. Sensitivity of the nf-core/airrflow and MiXCR pipelines on the data simulated with UMIs. Sensitivity was calculated for exact VDJ sequence matches to the truth repertoires (**A**), and matches that contain N nucleotides due to not reaching sufficient consensus (**B**). **C**. Number of incorrect sequences. **D**. Number of sequences present in the truth repertoires that were not identified by the pipelines. **E**. Number of clones identified by each pipeline. The discontinuous line indicates the true number of clones in the simulated repertoires. **F**. Mean clonal abundance (solid line) with max and min intervals (shaded area) from n=200 bootstrap samples of N=15,250 sequences of the three simulated repertoires.

In addition to recovering the original sequences in the sample, AIRR-seq data analysis often involves determining the clonal relationships of the individual sequences. This is important in order to assess whether a sequence comes from an expanded clone of the same progenitor cell. This step is particularly important in BCR data analysis, as mutations are introduced during clonal expansion by targeted somatic hypermutation. Thus, we assessed the ability of the workflow to recover the original number and size distribution of B-cell clones (Figure 2E). Close to 5,500 clones and 5,700 clones were identified for repA and repB (ground truth 5,100 for both), and 6,500 clones for repC (ground truth 6,305). The number of clones was overestimated by 7%, 11%, and 3%, respectively, but was robust with respect to simulated sequencing errors. The clonal abundance distribution (Figure 2F) reflected the true clonal distribution in all three repertoires for the simulated protocol with UMIs. Clonal inference by the Immcantation tools incorporated into nf-core/airrflow was superior to the inference method implemented in MiXCR (Figure 2E-F). The inferred number of clones and clonal abundance by both tools were affected by the increasing sequencing errors in the sans-UMI protocol, highlighting once more the importance of UMI error correction (Suppl. Figure 2 D and E).

### Case study: validating convergent antibody responses from public repositories

Previous studies have identified convergent antibody responses to the SARS-CoV-2 virus(8,44,61–63). These are sequences with high sequence similarity across patients, that can indicate BCRs targeting epitopes important for the neutralization of the SARS-CoV-2 virus or that are conserved across different strains. Robbiani *et al*.(44) identified a convergent antibody cluster comprising six BCR sequences across three COVID-19 infected subjects that were experimentally validated to bind the SARS-CoV-2 spike protein receptor binding domain (RBD). The identified convergent antibodies shared the same IGHV and IGHJ genes (IGHV1-58 and IGHJ3) and showed high similarity of the heavy chain CDR3 sequences (1 to 3 amino acid edits). To validate this finding and extend it to a larger number of subjects, we retrieved BCR repertoire datasets from COVID-19 diagnosed subjects from the AIRR Data Commons(45) by querying the iReceptor Gateway(46), together with healthy controls. These comprised datasets from 17 studies, including 496 antibody repertoires from 213 subjects sequenced with single-cell and bulk-based protocols, but did not include those reported by Robbiani *et al*. (Supl. Table 3). We processed the repertoires with nf-core/airrflow on an HPC cluster in just over 2 days (Figure 3B). 467 antibody repertoires from 196 subjects (99 healthy controls and 97 COVID-19 diagnosed) passed the QC criteria (Figure 3A, Suppl. Figure 3). We defined convergent antibody clusters by partitioning the processed sequences according to IGHV, IGHJ genes and junction length. We performed single-linkage hierarchical clustering on the junction amino acid sequences according to their pairwise hamming distances and defined convergent groups setting a length-normalized hamming distance threshold of 0.2, for each IGHV gene. We excluded convergent clusters that contained antibody sequences from a single subject, and clusters with sequences from healthy controls. We obtained 416 convergent antibody clusters of varying sizes with antibodies using the IGHV1-58 gene segment (Supplementary Figure 4, Figure3C). The convergent cluster with most sequences contained 1,189 sequences (182 unique junction sequences) from 20 subjects and 6 different studies (cluster 11312). This convergent cluster comprised BCRs with IGHV1-58 and IGHJ3 genes, and junction sequences at an edit distance of 1 to 3 amino acids from the six antibodies described by Robbiani *et al*. None of the other convergent clusters with IGHV1-58 and same junction length contained junction sequences at an edit distance of 3 amino acids or less. A sequence logo of all of the sequences in the convergent cluster 11312 revealed amino acid residue and properties conservation similar to the sequences reported in the original publication, with two conserved cysteines at positions 6 and 11, as well as two aspartate negative charges at positions 13 and 16 and a Phenylalanine at position 15 (Figure 3D and E), providing an indication of conserved specificity and the likely amino-acids important for binding. Thus, we conclude that this cluster represents the same convergent antibody sequences found by Robbiani *et al*., and extended this convergence to additionally 20 individuals. This use case highlights the applicability of nf-core/airrflow to validate findings originally made in small in-house cohorts with reanalysis of large publicly available AIRR-seq datasets.

**Figure 3.**
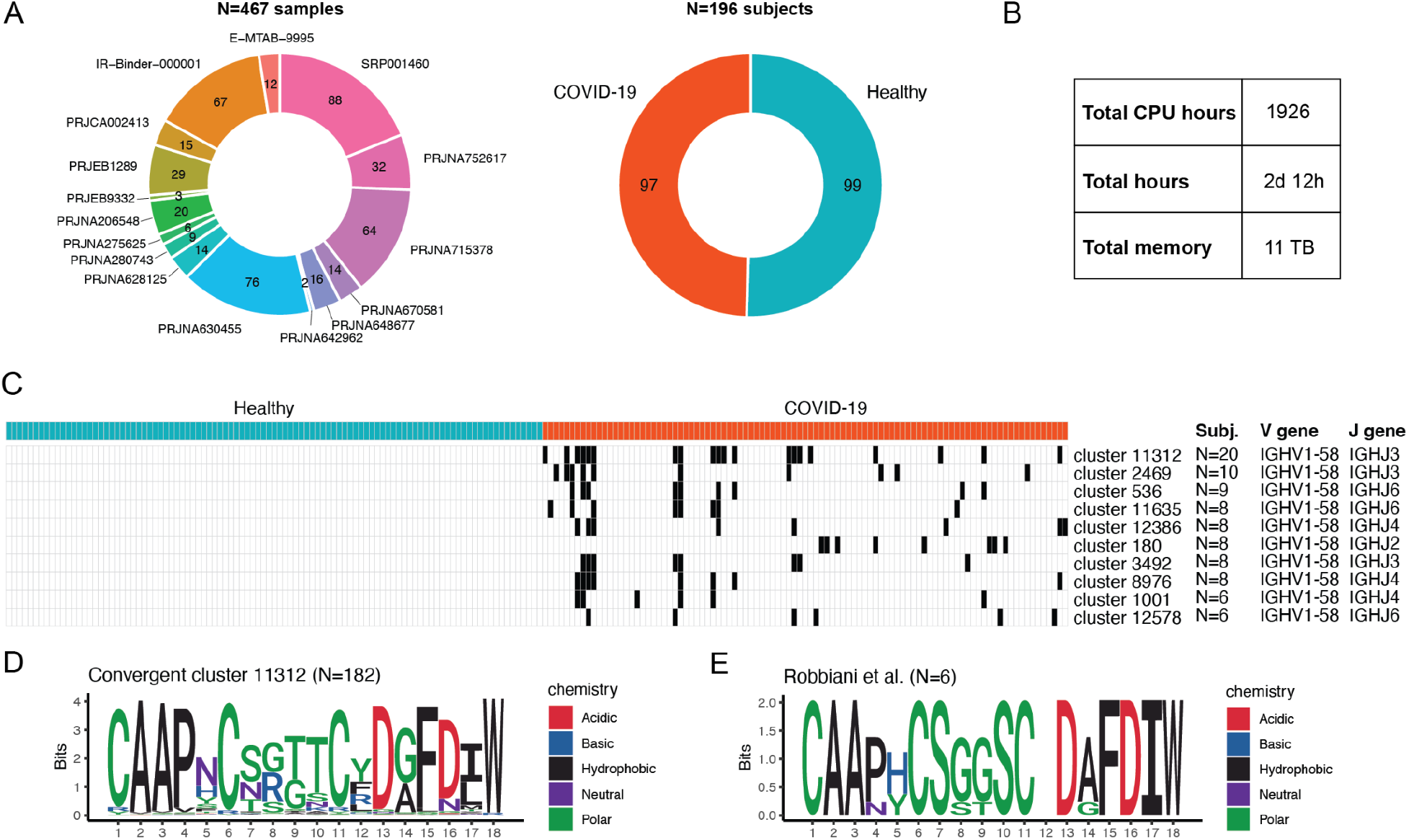
Application of nf-core/airrflow to process repertoires of COVID-19 infected or vaccinated participants together with healthy controls. **A**. Number of samples that passed the quality control after processing the data with nf-core/airrflow v3.2.0 and their distribution across projects. Number of subjects in each group (healthy or COVID-19 diagnosed). **B**. Resources needed to process the datasets on an HPC cluster. **C**. Top 10 antibody convergent clusters (rows) using the IGHV1-58 gene segment according to the number of unique COVID-19 subjects included in each cluster. Clusters containing sequences from a single individual, or from healthy controls were excluded. Each column represents a subject colored by their status (healthy or COVID-19 diagnosed). Black denotes the presence of sequences belonging to an antibody cluster for each particular subject. The annotations on the right denote the number of subjects with antibody sequences included in each convergent cluster, the heavy chain V gene assignment and the heavy chain J gene assignment of the majority of the sequences in the convergent cluster. **D**. Sequence logo of the N=182 junction sequences contained in the convergent cluster 11312. Coloring according to amino acid side chain properties. **E**. Sequence logo of the sequences published by Robbiani *et al*., positively tested against the SARS-CoV-2 spike protein receptor binding domain.

## Availability and future directions

We implemented nf-core/airrflow, a high-throughput, easy installation and portable workflow for the analysis of bulk and single-cell AIRR sequencing data. The workflow is available free of charge under the MIT license in GitHub (https://github.com/nf-core/airrflow), as part of the nf-core project. Detailed documentation and example results are available at https://nf-co.re/airrflow. The code to reproduce the data simulation, pipeline benchmark, and scripts to reproduce the analysis of the COVID and healthy datasets can be found on (https://bitbucket.org/kleinstein/projects).

We plan to further develop the workflow to incorporate additional analysis needs of the users. nf-core/airrflow current or future users are encouraged to join the nf-core community dedicated slack channel (https://nf-co.re/join) for questions or feature requests. As part of the nf-core community project, contributions to the workflow from other developers are welcome and encouraged, and will be reviewed before incorporation into the pipeline.

### Funding

This work was funded in part by the Chan Zuckeberg Initiative EOSS4 [2021-237742] to GG, the Deutsche Forschungsgemeinschaft (DFG) under Germany’s excellence Strategy [EXC 2180–390900677] (iFIT) and under [EXC 2124–390838134] (CMFI) to SN, and the NIH grant R01AI104739 to SHK.

### Disclosures

SHK receives consulting fees from Peraton. AP is an employee of Boehringer Ingelheim Pharma GmbH & Co KG and declares no conflict of interest. MCK has served on advisory boards and received speaker fees / travel grants from Merck, Sanofi-Genzyme, Novartis, Biogen, Janssen, Alexion, Celgene / Bristol-Myers Squibb and Roche. He has received research grants from Merck, Roche, Novartis, Sanofi-Genzyme and Celgene / Bristol-Myers Squibb. All other authors declare no conflicts of interest.

