## Supplementary information for "nf-core/airrflow: an adaptive immune receptor repertoire analysis workflow employing the Immcantation framework"

[1]Department of Pathology, Yale School of Medicine, New Haven, CT, USA.
[2]Quantitative Biology Center, Eberhard-Karls University of Tübingen, Tübingen, Germany.
[3]Yale Center for Research Computing, New Haven, CT, USA.
[4]Boehringer Ingelheim Pharma GmbH & Co. KG, Biberach, Germany.
[5]Program in Computational Biology and Bioinformatics, Yale University, New Haven, CT, USA.
[6]oNKo-Innate Pty Ltd, Melbourne, Victoria, Australia.
[7]Department of Computer Science, Eberhard-Karls University of Tübingen, Tübingen, Germany.
[8]M3 Research Center, University Hospital, Tübingen, Germany.
[9]Faculty of Engineering, Bar Ilan University, Ramat Gan, Israel.
[10]Department of Neurology and Stroke, Center for Neurology, Eberhard-Karls University of Tübingen, Tübingen, Germany.
[11]Hertie Institute for Clinical Brain Research, Eberhard-Karls University of Tübingen, Tübingen, Germany.
[12]Institute for Bioinformatics and Medical Informatics (IBMI), Eberhard-Karls University of Tübingen, Tübingen, Germany.
[13]Department of Immunobiology, Yale School of Medicine, New Haven, CT, USA.
*Equal contribution; [||]co-senior authors.

**Supplementary information** 1
Supplementary methods 2
Launching nf-core/airrflow 2
Processing full-size example data 2
Simulation of BCR sequencing data 3
Network analysis and plotting of simulated repertoires 4
Processing the BCR simulated datasets 4
Processing the simulated data with nf-core/airrflow 4
Processing the simulated data with MiXCR 5
Performance assessment 5
Processing publicly available COVID-19 datasets 5
References 6
Supplementary figures 9
Supplementary tables 13

### Supplementary methods

#### Launching nf-core/airrflow

Running nf-core/airrflow requires a Java (version ≥ 11) and Nextflow (version ≥ 23.05.0) installation. Additionally, a software manager such as Anaconda or a container engine such as Docker or Apptainer should be installed. The use of container engines is preferred over anaconda, to ensure reproducibility of the results across computing infrastructures. When launched, the pipeline will pull the containers that provide the software needed for each analysis step, to ensure reproducibility of the results. The full pipeline documentation with installation instructions can be found at https://nf-co.re/airrflow.

#### Processing full-size example data

To generate example results of the nf-core/airrflow v3.2.0 pipeline run, and verify that the pipeline runs on full-size datasets and on cloud infrastructures, we provide full-size test datasets extracted from the publication by Stern *et al.*(1). The raw fastq files were retrieved from the sequence read archive (SRA) with the accession number PRJNA248475. The dataset was processed on AWS batch using Nextflow Tower (https://tower.nf ) with the following command:

```
nextflow run nf-core/airrflow -r 3.2.0 -profile test_full \
--outdir s3://nf-core-awsmegatests/airrflow/results-333c8b521d62f6c61c658823d8831df93bc5b532 \
```

```
-w s3://nf-core-awsmegatests/work/airrflow/work-333c8b521d62f6c61c658823d8831df93bc5b532
```

The data was processed within 45 minutes and 16 seconds, utilizing 17.6 CPU hours. This test run is automatically triggered for each pipeline release on GitHub to ensure that each release runs without issues with full-size test data on cloud infrastructure. The full example results on this dataset processed can be explored on the nf-core website (https://nf-co.re/airrflow) at https://nf-co.re/airrflow/3.2.0/results/airrflow/results-333c8b521d62f6c61c658823d8831df93bc5b532.

#### Simulation of BCR sequencing data

**Simulating the V(D)J gene rearrangements**: 5,100 unique V(D)J rearrangements were generated with ImmuneSIM(2), which simulates the somatic recombination process occurring to generate full-length V(D)J sequences. Only heavy-chain sequences were considered for this analysis. The default V-, D- and J-gene usage distribution provided within immuneSIM was employed. For repertoires repA and repB, 100 sequences were simulated to undergo clonal expansion, whereas 5000 sequences were left unexpanded to simulate “singleton” sequences observed in real BCR repertoires extracted from PBMCs (Ruschil *et al.*(3)) that are important for the determination of clonal thresholds using hamming distance metrics. For repertoire C, the originally determined number of sequences in the same real BCR repertoire from with clonal expansion (1609) and singletons (4980) was maintained.

**Simulating clonal expansion**: to simulate clonal expansion and somatic hypermutation (SHM), lineage trees were simulated to resemble real BCR lineage trees (repA and repB) or extracted from a real BCR repertoire (CLAD1 baseline from Ruschil et al.(3)) which served as a reference sample (repC). The simulated lineage tree topologies were obtained with the *igraph v1.4.2 make_tree* function by providing the total number of nodes in the tree (the number of B-cells in the clone), the number of children per node and edge graph weights, which determines the number of mutations (hamming distance) from the parent to the children sequence. The number of individual nodes for each of the B-cell trees was set to follow a power-law distribution with alpha=2 (repA) or a uniform distribution (repB). The number of children nodes connected to each parent node was randomly chosen within a range from 2 to 11, the most frequent number of children per node observed in a reference BCR repertoire (CLAD1 baseline from Ruschil et al.) and maintained constant within a B-cell clone. The number of mutations from a parent to a children sequence was chosen between 1 and 40, following the probability distribution analyzed in the reference BCR repertoire from Ruschil et al. (Suppl. Figure 2). The lineage tree topologies were then populated with BCR sequences, setting as germline sequence the simulated ImmuneSIM rearrangements. For each B-cell lineage tree, somatic hypermutation was simulated with the *shmulateTree* function from SHazaM(4), providing the germline BCR sequence as input. The Human Heavy chain, Silent, 5-mer (HH-S5F) functional targeting model(4) was applied to introduce mutations to the germline sequences following the clonal lineage tree structure. The IGHM*01 sequence from IMGT was added 3’ of all VDJ simulated sequences.

**Simulating the library preparation method**: library preparation for the BCR repertoire was simulated using a 5' RACE strategy similar to the SMARTer TAKARA Bio protocol (https://www.takarabio.com/products/next-generation-sequencing/immune-profiling/human-reperB-celltoire/human-bcr-profiling-kit-for-illumina-sequencing). A dummy linker and spacer sequence were added to the 5' end of the VDJ sequences.
**Simulating the amplicon sequencing reads**: MiSeq sequencing read simulation was performed with Grinder v0.5.4(5), an amplicon sequencing read simulator. The C-region primer and linker + spacer sequences were provided for the targeted amplification simulation. Paired-end reads were simulated with a read length of 300 nt, and an insert distance following a normal distribution around 572 with a standard deviation of 5. Reads were simulated to reach an average 10-fold coverage per sequence. Read simulation was performed either with no introduced sequencing errors (0%) or with simulated sequencing errors following a linear distribution starting at 0.0001% at the beginning of reads and linearly increasing towards the end of the reads, with a value of 0.1%, 0.25%, 0.5% and 1.0% at the middle of the reads, respectively. These errors were set in accordance with previous studies on MiSeq sequencing error values and their distribution along the read positions(6,7).

**Simulating UMI-barcoded reads**: 12 nt long random sequences were added 5' of the R2 reads prior to amplification simulation with Grinder to simulate UMI-barcoded reads. Then the amplicon sequencing read simulation was performed as described above.

Suppl. Table 2 contains a list of the simulated repertoires and their characteristics. The simulated repertoire sequencing data files (fastq.gz) were uploaded to Zenodo (https://doi.org/10.5281/zenodo.10070476)

#### Network analysis and plotting of simulated repertoires

The synthetic B-cell clones generated with the igraph library (repA and repB) or reconstructed from real BCR repertoires analyzed with nf-core/airrflow were loaded into Cytoscape v3.9.1 for plotting the repertoires and performing network analysis to extract the edge weight distribution.

#### Processing the BCR simulated datasets

##### Processing the simulated data with nf-core/airrflow

The BCR simulated datasets were processed with the nf-core/airrflow v3.2.0 run with Nextflow v23.04.5.5708 and Java OpenJDK v17.0.1. The Singularity v1.8.7 container engine was employed. The pipeline was run on an HPC cluster with CentOS Linux 7 and a Slurm job scheduler (Slurm v21.08.8).
The launch script together with the metadata file and other inputs needed to run the pipeline can be found on the GitHub repository.

##### Processing the simulated data with MiXCR

The BCR simulated datasets were processed with MiXCR v4.3.2 on a desktop computer with 24 cores and 164 GB of memory running Ubuntu 22.04.3. The simulated sequencing data with the UMI protocol were processed with the *mixcr analyze generic-bcr-amplicon-umi* command whereas the data simulated with the sans-UMI protocol were analyzed with the *mixcr analyze generic-bcr-amplicon* command. The *mixcr findAlleles* command was then used to find the V(D)J alleles, followed by the mixcr findShmTrees command to reconstruct the B-cell clonal groups. The scripts to run mixcr for each repertoire and all the set parameters can be found on the GitHub repository.

#### Performance assessment

The following parameters were calculated for performance assessment:

- **Sensitivity exact matches**: proportion of all sequences in the ground truth repertoire that were correctly identified by the pipeline.

$$Sensitivity = \frac{exact\ matches}{all\ sequences\ in\ truth\ set}$$

- **Sensitivity matches with N nucleotides**: proportion of all sequences in the ground truth repertoire that were correctly identified by the pipeline, including sequences that contained N nucleotides (insufficient consensus to assign the base).

$$Sensitivity\ matches\ with\ Ns = \frac{exact\ matches + matches\ with\ Ns}{all\ sequences\ in\ truth\ set}$$

#### Processing publicly available COVID-19 datasets

iReceptor was queried for publicly available datasets on COVID-19 diagnosed individuals and healthy controls on September 12, 2023. The query for COVID-19 diagnosed subjects included the terms: diagnosis “COVID-19”, study group “case” and PCR target “IGH, IGK or IGL”. The search returned 707 repertoires from 12 studies. 2 studies were eliminated (IR-Roche-000001, and PRJNA638224) as the BCR sequences were not available on the database, leaving 10 COVID-19 studies (N=289 repertoires, N=105 subjects). Two of these studies contained healthy controls. To retrieve additional repertoires from healthy controls, a second query was performed with the terms: study group “control” and PCR target “IGH”, “IGK” or “IGL”. The search returned 409 repertoires from 10 studies, all submitted prior to 2018. One study was eliminated as the BCR sequences were not available on the database (IR-Roche-000001), and the 9 remaining studies were included in the analysis as Healthy subjects (N=207 repertoires, N=108 subjects) (**Supplementary Table 4**).

The datasets were processed using nf-core/airrflow v3.2.0 on an HPC cluster running SLURM limited to 2 nodes with 36 CPUs/node and 163 GB of memory/node (Intel Xeon Gold 6240 central computing unit processor). The command used to process the datasets and nextflow configuration file can be found on the Github repository. After processing with nf-core/airrflow a total of 467 samples (289 samples from 97 COVID-19 individuals, and 178 samples from 99 healthy individuals) passed the quality control steps and were included in the convergent antibody analysis (**Supplementary Figure 3**).

### Supplementary figures

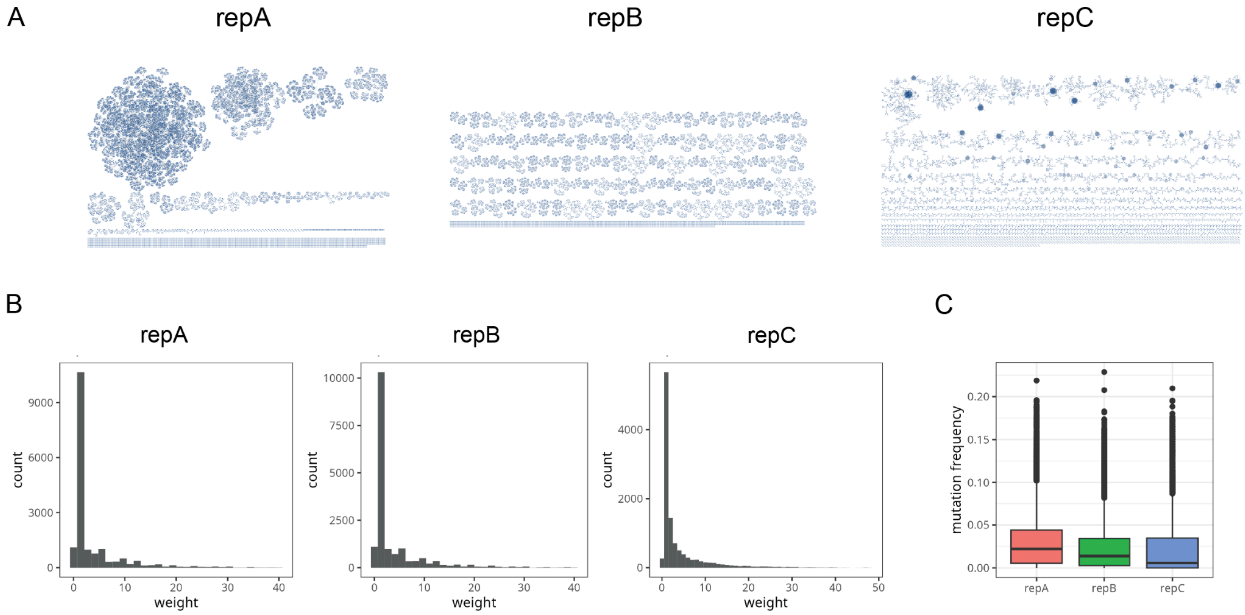


**Supplementary Figure 1. Simulated repertoires for pipeline benchmarking. A.** Network visualization of the simulated repertoires. ImmuneSIM was used to simulate somatic recombination whereas the SHazaM *shmulateTree* function was used to simulate somatic hypermutation. Two simulated repertoires were generated containing 100 B-cell clones and 5000 non-clonally expanded singletons each: a repertoire with clonal abundances following a power-law distribution with *alpha*=2 (repA), and a repertoire with uniform clonal abundances (repB). A third repertoire was simulated utilizing clonal trees recovered from a real BCR repertoire sample (repC). Sequencing data was simulated with unique molecular identifiers addition (UMI), and without (sans-UMI). BioGrinder was used to generate the simulated amplicon sequencing fastq files with increasing percentages of simulated sequencing errors (0 - 1.0% in the middle of reads). **B.** Histogram of the lineage tree edge weights, representing the number of mutations from a parent sequence to a child sequence in the lineage tree. **C.** Mutation frequencies of all sequences in repA, repB and repC.

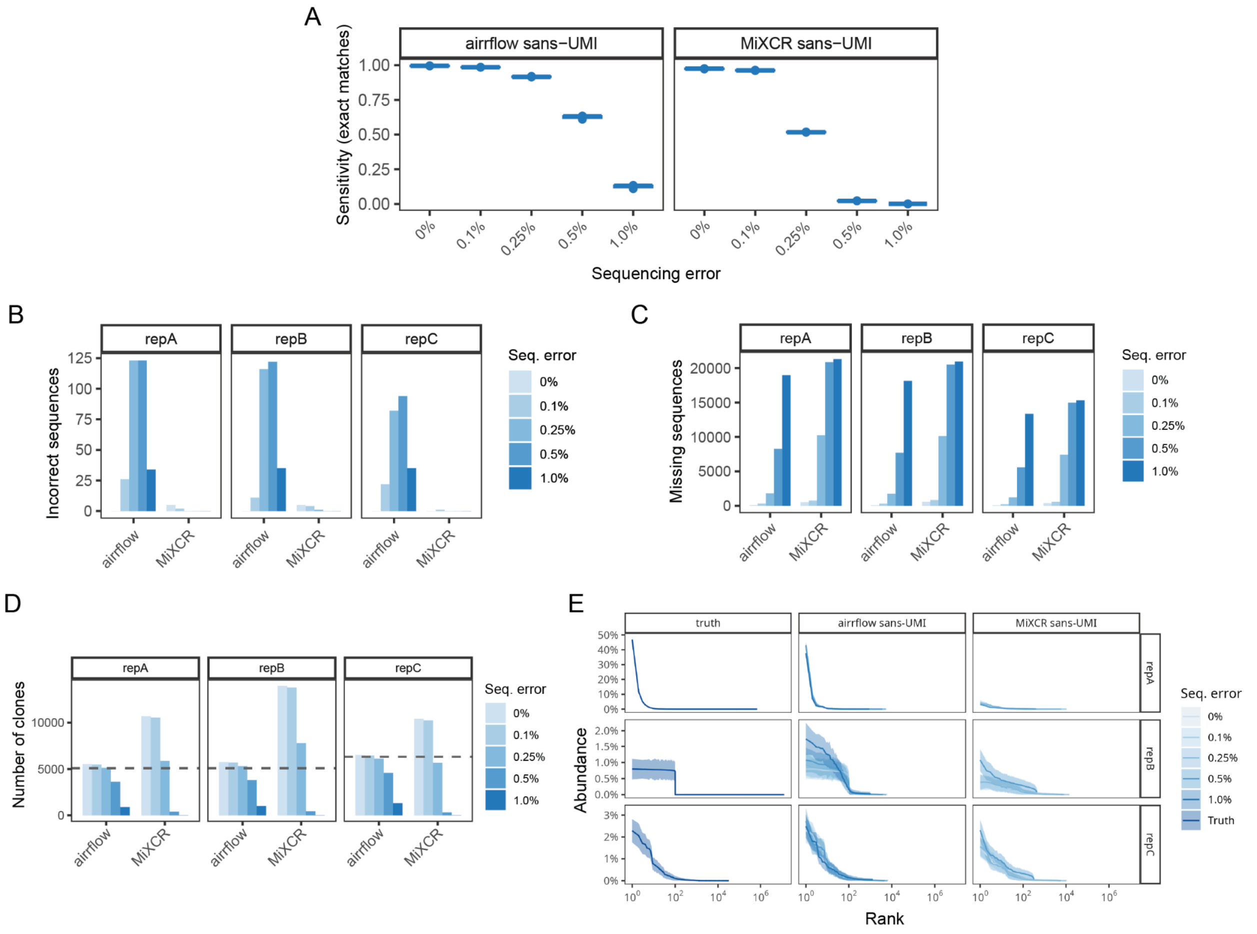


**Supplementary Figure 2. Performance evaluation of the nf-core/airrflow pipeline on synthetically generated BCR repertoires for the sans-UMI protocol**. Sensitivity (A), number of incorrect sequences (B) and number of missing sequences (C) for each of the simulated repertoires with increasing sequencing errors. Number of clones (D) and clonal abundance (E) for each of the repertoires. The horizontal gray discontinuous line indicates the true number of simulated clones.

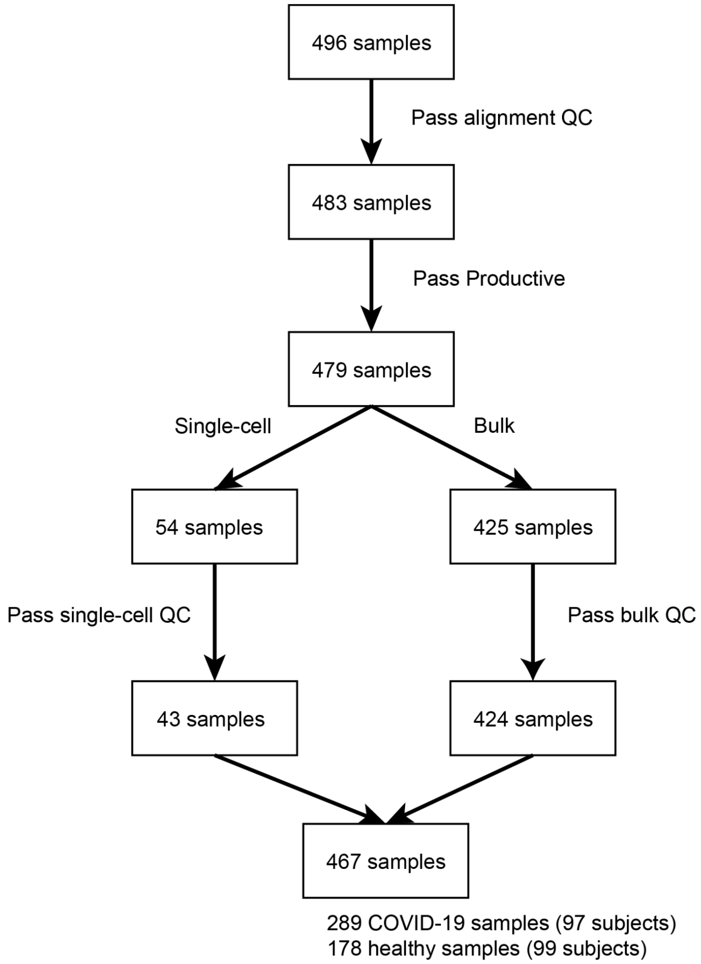


**Supplementary Figure 3. Processing of 496 samples from COVID-19 diagnosed and healthy control subject with nf-core/airrflow v3.2.0.** The diagram shows the number of samples remaining after each pipeline QC steps. 17 samples were eliminated as they did not have any remaining sequences that passed the alignment QC (more than 200 informative positions, less than 10% N nucleotides, productive VDJ sequences. One sample did not pass the bulk QC criteria. Eleven samples were eliminated during the single-cell QC analysis, as they did not have any remaining cells after filtering out cells with only light chains, cells with multiple heavy chains, and spurious sequences with the same cell barcode and exact same BCR VDJ sequence.

A

B

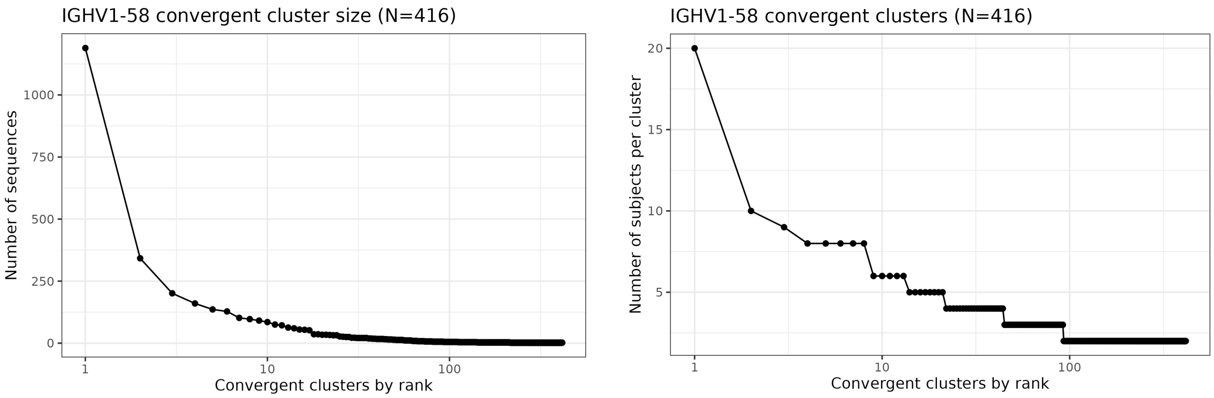


**Supplementary Figure 4. Convergent clusters using the IGHV1-58 gene segment.** Convergent clusters with sequences from a single individual orcontaining any sequences from healthy subjects were excluded. **A.** Size distribution of the convergent clusters ranked by size. **B.** Number of subjects per cluster for the convergent clusters ranked by number of subjects in each cluster.

### Supplementary tables

**Supplementary table 1. Comparison of functionality between the nf-core/airrflow v3.2.0 and the MiXCR v4.3.2 pipeline.**

| Functionality | | nf-core/airrflow v3.2.0 (Immcantation) | MiXCR v4.3.2 |
|---|---|---|---|
| **Supported protocols** | **Bulk** | yes | yes |
| | **Single-cell** | yes | yes |
| | **Mixed bulk and single-cell** | yes | no |
| **Read assembly** | **UMI barcode support** | yes | yes |
| **V(D)J & C reference alignment** | **Alignment tool** | IgBLAST | Own algorithm |
| **Quality control** | **Detection of cross-sample contamination** | yes | no |
| **Clonal analysis** | **Define clones** | yes | yes |
| | **Allows specification of clonal threshold** | yes | no |
| | **Lineage tree reconstruction** | yes | yes |
| **Repertoire analysis** | | yes | yes |
| **Usability** | **Reproducibility** | yes | When using container |
| | **One-command execution** | yes | no |
| | **Parallelization across samples** | yes | no |
| | **Portability to cloud** | yes | limited |

**Supplementary Table 2**. **BCR repertoire sequencing simulation runs.** Run identifier (Run ID), original repertoire sequences (Repertoire), the addition of UMIsin the library simulation procedure (UMI), simulated sequencing error percentage at the center of the reads (Sequencing error).

| Run ID | Repertoire | UMI | Sequencing error |
|---|---|---|---|
| rep000 | repA | no | 0% |
| rep001 | repB | no | 0% |
| rep002 | repA | no | 0.1% |
| rep003 | repA | no | 0.25% |
| rep004 | repA | no | 0.5% |
| rep005 | repA | no | 1.0% |
| rep006 | repB | no | 0.1% |
| rep007 | repB | no | 0.25% |
| rep008 | repB | no | 0.5% |
| rep009 | repB | no | 1.0% |
| rep010 | repA | yes | 0% |
| rep011 | repB | yes | 0% |
| rep012 | repA | yes | 0.1% |
| rep013 | repA | yes | 0.25% |
| rep014 | repA | yes | 0.5% |
| rep015 | repA | yes | 1.0% |
| rep016 | repB | yes | 0.1% |
| rep017 | repB | yes | 0.25% |
| rep018 | repB | yes | 0.5% |
| rep019 | repB | yes | 1.0% |
| rep101 | repC | no | 0% |
| rep102 | repC | no | 0.1% |

| | | | |
|---|---|---|---|
| rep103 | repC | no | 0.25% |
| rep104 | repC | no | 0.5% |
| rep105 | repC | no | 1.0% |
| rep111 | repC | yes | 0% |
| rep112 | repC | yes | 0.1% |
| rep113 | repC | yes | 0.25% |
| rep114 | repC | yes | 0.5% |
| rep115 | repC | yes | 1.0% |

**Supplementary Table 3**. **Studies, subjects and repertoires included in the healthy vs COVID-19 infected participants dataset.**

| Study ID | Mode | Number of subjects | Number of repertoires | Status | Reference |
|---|---|---|---|---|---|
| E-MTAB-9995 | Single-cell | 4 | 12 | COVID-19 | Sokal et al. 2021(8) |
| IR-Binder-000001 | Bulk | 35 | 67 | COVID-19 | Schultheiss et al. 2020(9) |
| PRJCA002413 | Single-cell | 15 | 15 | COVID-19, Healthy | Wen et al. 2020(10) |
| PRJNA628125 | Bulk | 7 | 14 | COVID-19 | Nielsen et al. 2020(11) |
| PRJNA630455 | Bulk | 13 | 76 | COVID-19, Healthy | Kuri-Cervantes et al. 2020(12) |
| PRJNA648677 | Bulk | 7 | 16 | COVID-19 | Kim et al. 2020(13) |
| PRJNA715378 | Bulk | 5 | 64 | COVID-19 | Schmitz et al. 2021(14) |
| PRJNA752617 | Bulk | 4 | 32 | COVID-19 | Goel et al. 2021(15) |
| PRJNA642962 | Single-cell | 1 | 2 | COVID-19 | Woodruff et al. 2020(16) |
| PRJNA670581 | Single-cell | 14 | 14 | COVID-19 | Mor et al. 2021(17) |
| SRP001460 | Bulk | 37 | 88 | Healthy | Boyd et al. 2009(18) |
| PRJEB9332 | Bulk | 8 | 20 | Healthy | Chang et al. 2016(19) |
| PRJEB1289 | Bulk | 29 | 29 | Healthy | Bashford-Rogers et al. 2013(20) |
| PRJNA206548 | Bulk | 7 | 20 | Healthy | Michaeli et al. 2014(21) |
| PRJNA275625 | Bulk | 6 | 6 | Healthy | Valdés-Alemán et al. 2014(22) |
| PRJNA280743 | Bulk | 10 | 10 | Healthy | Tipton et al. 2015(23) |
| PRJNA381394 | Single-cell | 11 | 11 | Healthy | Vergani et al. 2017(24) |
| **TOTAL** | | 213 | 496 | | |

**Supplementary Table 4**. **Top 10 convergent clones found across COVID-19 infected subjects, and not in healthy subjects, with the IGHV1-58 gene**. The J call column indicates the J call of the majority of the sequences in the convergent group. The junction nucleotide sequence length is also indicated. The number of subjects, samples, and studies included in each convergent group.

| **Convergent group** | **V call** | **J call** | **Junction length (nt)** | **# subjects** | **# samples** | **# studies** |
|---|---|---|---|---|---|---|
| group 11312 | IGHV1-58 | IGHJ3 | 54 | 20 | 24 | 6 |
| group 2469 | IGHV1-58 | IGHJ3 | 39 | 10 | 14 | 5 |
| group 536 | IGHV1-58 | IGHJ6 | 30 | 9 | 10 | 4 |
| group 11624 | IGHV1-58 | IGHJ6 | 57 | 8 | 12 | 3 |
| group 12386 | IGHV1-58 | IGHJ4 | 57 | 8 | 10 | 5 |
| group 180 | IGHV1-58 | IGHJ2 | 63 | 8 | 8 | 1 |
| group 3489 | IGHV1-58 | IGHJ3 | 42 | 8 | 8 | 3 |
| group 8976 | IGHV1-58 | IGHJ4 | 51 | 8 | 11 | 4 |
| group 1001 | IGHV1-58 | IGHJ4 | 36 | 6 | 6 | 4 |
| group 12578 | IGHV1-58 | IGHJ6 | 57 | 6 | 6 | 5 |
